# Pits and CtBP control tissue growth in *Drosophila melanogaster* with the Hippo pathway transcription repressor, Tgi

**DOI:** 10.1101/847707

**Authors:** Joseph H.A. Vissers, Lucas G. Dent, Colin House, Shu Kondo, Kieran F. Harvey

**Affiliations:** Peter MacCallum Cancer Centre, 305 Grattan Street, Melbourne, Victoria, Australia, 3000; Sir Peter MacCallum Department of Oncology, University of Melbourne, Parkville, Victoria, Australia, 3010; Laboratory of Invertebrate Genetics, National Institute of Genetics, 1111 Yata, Mishima, Shizuoka, Japan; Department of Anatomy and Developmental Biology, and Biomedicine Discovery Institute, Monash University, Clayton, Australia, 3800

**Keywords:** Hippo pathway, tissue growth, transcription, *Drosophila*

## Abstract

The Hippo pathway is an evolutionary conserved signalling network that regulates organ size, cell fate control and tumorigenesis. In the context of organ size control, the pathway incorporates a large variety of cellular cues such as cell polarity and adhesion into an integrated transcriptional response. The central Hippo signalling effector is the transcriptional co-activator Yorkie, which controls gene expression in partnership with different transcription factors, most notably Scalloped. When it is not activated by Yorkie, Scalloped can act as a repressor of transcription, at least in part due to its interaction with the corepressor protein Tgi. The mechanism by which Tgi represses transcription is incompletely understood and therefore we sought to identify proteins that potentially operate together with it. Using an affinity purification and mass-spectrometry approach we identified Pits and CtBP as Tgi-interacting proteins, both of which have been linked to transcriptional repression. Both Pits and CtBP were required for Tgi to suppress the growth of the *D. melanogaster* eye and *CtBP* loss suppressed the undergrowth of *yorkie* mutant eye tissue. Furthermore, as reported previously for Tgi, overexpression of Pits suppressed transcription of Hippo pathway target genes. These findings suggest that Tgi might operate together with Pits and CtBP to repress transcription of genes that normally promote tissue growth. The human orthologues of Tgi, CtBP and Pits (VGLL4, CTBP2 and IRF2BP2) physically and functionally interact to control transcription, implying that the mechanism by which these proteins control transcriptional repression is conserved throughout evolution.

## INTRODUCTION

The Hippo pathway was first discovered in *D. melanogaster* genetic screens as an important regulator of organ growth (Kango-Singh *et al*. 2002; Tapon *et al*. 2002; Harvey *et al*. 2003; Jia *et al*. 2003; Pantalacci *et al*. 2003; Udan *et al*. 2003; Wu *et al*. 2003). It has subsequently been shown to control the growth of multiple different tissues – epithelial, muscle, neural and blood – in different species (Halder and Johnson 2011). It also controls cell fate choices in both *D. melanogaster* and mammals, whilst mutation of Hippo pathway genes underpins several human cancers (Harvey *et al*. 2013). In the context of organ size control, the Hippo pathway responds to cell biological cues and its surrounding environment. For example, the Hippo pathway is regulated by cell-cell adhesion, cell-matrix contacts, cell polarity proteins and by mechanical forces transmitted by the actin and spectrin cytoskeletons (Halder *et al*. 2012; Gaspar and Tapon 2014; Zheng and Pan 2019). These signals converge on a core signalling complex consisting of the Hippo (Hpo) and Warts (Wts) kinases and the adaptor proteins Salvador (Sav) and Mats (Irvine 2012). The central Hippo signalling effector is Yorkie (Yki), which is a transcriptional co-activator. Yki rapidly shuttles between the nucleus and cytoplasm and this is regulated by Wts-mediated phosphorylation at conserved amino acids, which limits access to the nucleus (Dong *et al*. 2007; Zhao *et al*. 2007; Oh and Irvine 2008; Manning *et al*. 2018). When nuclear, Yki partners with sequence specific transcription factors to control expression of genes such as *DIAP1*, *bantam* and *cyclin E*.

The best-characterized Yki-interacting transcription factor is the TEA domain protein Scalloped (Sd) (Goulev *et al*. 2008; Wu *et al*. 2008; Zhang *et al*. 2008). Yki and Sd promote transcription by interacting with chromatin modifying proteins like the Mediator complex, the SWI-SNF complex and the Trithorax-related histone methyltransferase complex (Zheng and Pan 2019). The mammalian orthologues of Yki (YAP and TAZ) and Sd (TEAD1-TEAD4) regulate transcription by interacting with similar protein complexes. In human cells, YAP and TEAD regulate gene expression predominantly by binding to enhancers, as opposed to promoters (Stein *et al*. 2015; Zanconato *et al*. 2015). In addition, they promote transcriptional elongation by recruiting the Mediator complex and Cdk9 (Galli *et al*. 2015).

The mechanism by which Sd and TEADs repress transcription is less well defined. Genetic evidence from tissues such as the *D. melanogaster* ovary indicates that Sd can act as a default repressor of transcription, and this activity is antagonized by Yki (Koontz *et al*. 2013). Sd’s default repressor function is mediated in part by the corepressor protein Tondu-domain-containing Growth Inhibitor (Tgi) (Koontz *et al*. 2013), also known as Sd-Binding-Protein (Guo *et al*. 2013). In order to activate gene expression, Yki is thought to compete with Tgi for Sd binding and alleviate repression of target genes. Tgi interacts via its Tondu domains with Sd and via PY motifs with Yki’s WW domains (Guo *et al*. 2013; Koontz *et al*. 2013). This relationship is conserved in mammals between YAP and TAZ, TEAD1-TEAD4 and VGLL4 (the Tgi orthologue) (Koontz *et al*. 2013; Zhang *et al*. 2014). Sd can also regulate transcription together with the Zinc finger domain protein Nerfin-1. These proteins work in partnership required to maintain the fate of *D. melanogaster* medulla neurons (Vissers *et al*. 2018) and they also influence cell competition in growing imaginal discs (Guo *et al*. 2019). Currently, the mechanism by which Tgi and VGLL4 cooperate with Sd/TEADs to control transcription and tissue growth is incompletely understood.

To better understand how Tgi regulates transcription we used proteomics approaches and identified four high confidence Tgi-interacting proteins, all of which are transcriptional regulatory proteins. In addition to the known Tgi partners Yki and Sd, we identified two previously unknown Tgi-interacting proteins, CG11138 (also known as protein interacting with Ttk69 and Sin3A, or Pits) and C-terminal binding protein (CtBP), both of which have been linked to transcriptional repression (Turner and Crossley 2001; Liaw 2016). Both gain and loss of function of *pits* and *CtBP* modified tissue growth aberrations caused by Hippo signalling defects. Furthermore, overexpression of Pits reduced expression of well-defined Hippo pathway target genes, thus highlighting the possibility that Pits and CtBP operate with Tgi to limit tissue growth and transcription.

## MATERIALS AND METHODS

### *D. melanogaster* strains

Transgenic *D. melanogaster* strains harbouring *UAS-pits* were generated by Bestgene (USA). For experiments on adult animals, the *UAS-pits* transgenic line 2M was used. For experiments on larval imaginal discs, the *UAS-pits* transgenic line 4M was used. Other stocks were *CtBP^87De-10^, CtBP^03463^, mCherry RNAi* (BSC#35785), *CtBP RNAi* (BSC#32889), *w^1118^ P(EP)pits^EP1313^, y^1^ P(SUPor-P)pits^KG07818^, en-Gal4, ptc-Gal4, y w P W*+ *[ubi-GFP] FRT19A; eyFlp, bantam-lacZ, ex^697^, UAS-lacZ, UAS-GFP, UAS-RFP, UAS-EYFP* (all Bloomington *Drosophila* Stock Centre), *CtBP RNAi KK107313, pits RNAi KK101998, β-galactosidase RNAi GD51446* (all Vienna *Drosophila* RNAi Centre), *sav^2^* (Tapon *et al*. 2002), *UAS-HA-Tgi* (Koontz *et al*. 2013) (a gift from DJ Pan). As has been reported for many VDRC KK RNAi lines (Vissers *et al*. 2016), the *CtBP* KK RNAi line contains two transgenic insertion sites. For this study, a recombinant strain was generated harbouring only a *CtBP* KK RNAi insertion site only at 30B.

### Generation of *pits* mutant alleles

Mutant alleles of *pits* were generated using the transgenic CRISPR/Cas9 system, as previously described (Kondo and Ueda 2013), in the *y w FRT19A* background. The 20-bp target sequence in the gRNA was GCCCAGCGAACTGAGCGAAC, which resides in the coding region of the first exon. After screening eight candidate mutant strains by sequencing, we obtained two alleles, *pits^SK2^* and *pits^SK5^*, that carry 17-bp and 8-bp deletions, respectively, and result in premature termination of translation due to frameshift mutations. The deletions in these *pits* alleles are indicated below:

WT: GTGGTCCAGGTCCGTCGGGCGCCGGTTCGCTCAGTTCGCTGGGCAATGCCGTTGC
SK2: GTGGTCCAGGTCCGTCGGGCGC-----------------TGGGCAATGCCGTTGC -17
SK5: GTGGTCCAGGTCCGTCGGGCGC--------TCAGTTCGCTGGGCAATGCCGTTGC -8

### Plasmids

The constructs encoding Tgi fused to Streptavidin Binding Protein (SBP) tag were cloned by PCR amplification of *tgi* transcript variant B from pAc-Myc-Tgi (a gift from DJ Pan). XhoI, SpeI digested PCR products were ligated in XhoI, SpeI digested pMK33-SBP-N and pMK33-SBP-C (Kyriakakis *et al*. 2008). The Tgi mutant constructs used for interaction domain mapping were synthesized (Biomatik) and subcloned into pMK33-SBP-C using SpeI and XhoI. *pUASt-Pits* was generated by synthesis of *pits* transcript variant C (NM_132581.3) flanked by BglII and XhoI sites (Biomatik) and subcloning into pUASt.

### S2 cell culture and transfection

S2 cells were maintained in Schneider’s medium supplemented with 10% FBS and 1% penicillin/streptomycin as in (Poon *et al*. 2018). For stable transfections, 3 million cells were seeded and transfected with 1μg pMK33-based plasmid DNA using Effectene (Qiagen). After 48 hours, cells were cultured in 300μg/ml hygromycin for approximately 1 month to kill untransfected cells, and subsequently cultured under continuous drug selection.

### Affinity Purification and Mass Spectrometry

Streptavidin Binding Protein tagged proteins were purified as performed previously (Veraksa *et al*. 2005; Degoutin *et al*. 2013), with minor modifications. Cells were induced with 75mM CuSO4 overnight, washed in ice-cold PBS twice and lysed in 50 mM Tris pH 7.5, 5 % glycerol, 0.2 % IGEPAL, 1.5 mM MgCl_2_, 125 mM NaCl, 25 mM NaF, 1 mM Na_3_VO_4_, 1 mM DTT and Complete protease inhibitors (Roche). Lysis was allowed to proceed for 20 minutes, after which lysates were cleared by centrifugation. At this point, an input sample was taken for western analysis and the remaining lysate was incubated with streptavidin beads (Pierce) for 4 hrs at 4°C. Beads were washed with lysis buffer 4 times. For mass spectrometry analysis, bound proteins were eluted using biotin. For western blot analysis, bound proteins were denatured by addition of 1 bed volume of 2x LDS buffer supplemented with 1x sample reducing agent (Life Technologies) and incubated at 70°C for 5 minutes.

### Western blot analysis

NuPage and Bolt 4-12% BisTris SDS-PAGE gels (Life Technologies) were run at 200V in 1x MOPS buffer supplemented with antioxidant. Proteins were transferred to methanol-activated Immobilon membrane (Millipore) in transfer buffer (Tris, Glycine) in ice 1.5 hrs at 110V. Membranes were blocked in 5% (w/v) powder milk dissolved in Tris buffered Saline with 0.1% Tween (TBS/T) at least 30 minutes at RT. Membranes were incubated with primary antibodies o/n at 4°C in blocking buffer, washed 3x 10 minutes in TBS/T, incubated with HRP-coupled secondary antibodies 45 minutes RT, and washed 3x 10 minutes in TBS/T. Membranes were incubated with ECL Prime (GE) and western blot data were detected using a BioRad Chemidoc system. Primary antibodies were specific to the SBP epitope tag (mouse sc-101595, Santa Cruz), CtBP (goat sc-26610, Santa Cruz), Pits (rabbit, a gift from V. Corces) and Tubulin (mouse, T5168, Sigma).

### Immunofluorescence

For staining of larval tissues and pupal eye discs, samples were dissected into 4% (v/v) paraformaldehyde (PFA)/PBS and fixed for 20 minutes. After fixation and staining, tissues were washed 3 times for 10 minutes each in PBT (PBS supplemented with 0.3% Triton X-100). Primary and secondary antibody incubations were overnight at 4°C in PBT (0.1%) with 10% Normal Goat Serum (NGS). Primary antibodies were specific to β-galactosidase (Sigma, mouse. 1:100), DIAP1 (a gift from B. Hay, mouse, 1:00), Pits (a gift from V. Corces, rabbit, 1:500), HA-epitope-tag (Roche, rat, 1:100), Discs Large (DSHB, mouse, 1:50). Secondary antibodies (1:500) were from Invitrogen. DAPI was used to visualize nuclei. Images were acquired on a Nikon C2 confocal microscope. Image analysis was performed in Fiji (Schindelin *et al*. 2012), Adobe Photoshop and Adobe Illustrator.

## RESULTS

### Tgi forms a physical complex with Pits and CtBP via conserved motifs

To identify Tgi-interacting proteins we generated S2 cell lines that stably expressed Copper-inducible Streptavidin Binding Protein (SBP)-tagged Tgi or SBP alone. To alleviate the potential that the SBP tag could affect recovery of Tgi-binding proteins, we generated both N- and C-terminal Tgi-SBP fusion constructs. SBP purifications were performed for control empty vector SBP and Tgi-SBP and analysed by mass spectrometry. We recovered only four proteins with high confidence in Tgi purifications, compared to controls: the known Tgi-interacting proteins Yki and Sd, as well as CtBP and Pits (Figure 1A). Interestingly, both CtBP and Pits have been defined as regulators of transcription and most commonly reported as transcriptional co-repressors (Turner and Crossley 2001; Liaw 2016). CtBP is part of the CoREST co-repressor complex that interacts with many transcription factors (Turner and Crossley 2001). Pits is a nuclear protein (Rohrbaugh *et al*. 2013), and contains an N-terminal zinc finger domain and a C-terminal RING domain connected by a long region of low complexity (Liaw 2016).

**Figure 1.**
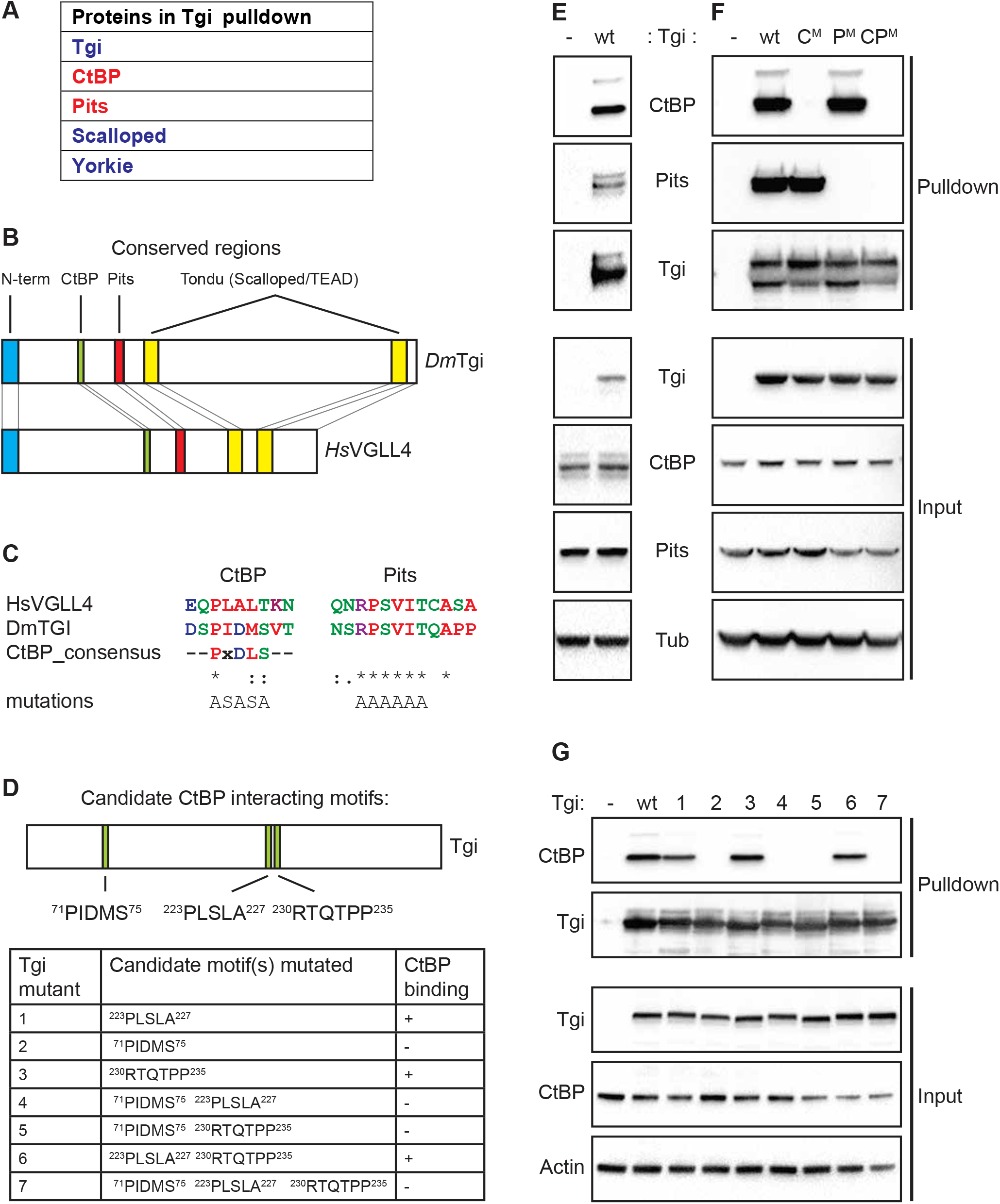
Tgi forms a physical complex with Pits and CtBP via conserved motifs. (A) Proteins identified by mass spectrometry with high confidence in Tgi-SBP purifications. (B) Schematic diagram of amino acid motifs conserved between *D. melanogaster* Tgi and human VGLL4. (C) Alignment of predicted CtBP and Pits interaction motifs in *D. melanogaster Tgi* (transcript variant B) and human *VGLL4* (isoform B). Alignment to the consensus CtBP interaction motif is shown for comparison. Four our biochemical studies, the PIDMS CtBP interaction motif in Tgi was mutated to ASASA and the Pits interaction motif was mutated to AAAAAA. (D) Schematic diagram of Tgi with the positions of candidate CtBP interaction motifs and a Table showing the amino acid sequence of these motifs. In the right column of the Table is a summary of CtBP and Tgi interaction experiments from (G). (E and F) Western blots of wildtype Tgi-SBP (E) and mutant (F) Tgi-SBP purifications and input lysates, using the indicated antibodies. C^M^, harbours a mutation in the predicted CtBP interaction motif in Tgi (PIDMS to ASASA), whilst P^M^, harbours a mutation in the predicted Pits interaction motif in Tgi (RPSVIT to AAAAAA). CP^M^ has mutations in both motifs. (G) Western blots of wildtype and mutant Tgi-SBP purifications and input lysates, using the indicated antibodies. Three candidate CtBP interaction motifs were mutated either singly, doubly or triply, as indicated in (D). Motif mutations were: PIDMS to ASASA, PLSLA to ASASA and RTQTPP to AAQTAA.

To verify our proteomics studies, we repeated the Tgi-SBP purifications and also designed structure-function studies to identify conserved motifs in Tgi that might mediate its interaction with CtBP and Pits. To determine the protein domains of Tgi responsible for interacting with CtBP and Pits, we compared the amino acid sequences of Tgi and VGLL4 to identify regions of evolutionary conservation. Tgi and VGLL4 contain two conserved Tondu domains responsible for the interaction with Sd and TEAD transcription factors, respectively (Figure 1B) (Guo *et al*. 2013; Koontz *et al*. 2013). In addition, as reported previously (Barrionuevo *et al*. 2014), we observed three conserved regions in Tgi: the N terminal region, and two short motifs between this region and the first Tondu domain (Figure 1B). The first short conserved Tgi motif (PIDMS) bears strong resemblance to a consensus CtBP-interaction motif (PxDLS, where x is any amino acid) (Figure 1C). We mutated this motif and two further potential CtBP interacting motifs [based on (Schaeper *et al*. 1995; Postigo and Dean 1999; Turner and Crossley 2001; Quinlan *et al*. 2006)] (Figure 1C and D). In mammals, a protein fragment encompassing the second short motif (RPSVIT) and the first Tondu domain of VGLL4 is sufficient to mediate the interaction with IRF2BP2 (Teng *et al*. 2010). As such, we also generated Tgi-SBP constructs bearing single and double mutations of the PIDMS and RPSVIT motifs (Figure 1C).

Initially, we subjected Tgi-SBP pulldowns to SDS-PAGE and probed for CtBP and Pits using antibodies to detect the endogenously expressed proteins. Western blot analysis verified both interactions and revealed two bands for CtBP, and three bands for Pits (Figure 1E). We repeated these pulldowns with both wild-type and mutants Tgi proteins and found that, as predicted, Tgi bound to CtBP via the PIDMS motif, and to Pits via the RPSVIT motif (Figure 1F). The importance of the Tgi PIDMS motif for interaction with CtBP was further confirmed by performing pulldowns with the three Tgi variants described in (Figure 1D). This experiment revealed that only mutation of the PIDMS motif prevented the interaction of Tgi with CtBP (Figure 1D and G). In mammals, physical interactions have been reported between the Tgi orthologue, VGLL4 and the three Pits orthologues (Interferon Regulatory Factor 2-Binding Protein Like, 1 and 2; IRF2BPL, IRF2BP1 and IRF2BP2, respectively) (Teng *et al*. 2010; Huttlin *et al*. 2015). A protein-protein interaction between the VGLL4-TEAD4 complex and CtBP orthologue CTBP2 has also been reported (Zhang *et al*. 2018). These studies provide independent validation of our Tgi-SBP purifications and indicate that the mode of interaction between Tgi, Pits and CtBP is conserved between *D. melanogaster* and mammals.

### CtBP limits eye growth and is required for Tgi-induced growth inhibition

Next, we investigated genetic interactions between *CtBP* and the Hippo pathway. We used the eyeless-Flp system in combination with the FRT cell lethal system to generate *sav* and *CtBP* single and double mutant heads, given that both genes are on chromosome arm 3R. As reported previously, generation of tissue homozygous mutant for the hypomorphic *sav^2^* allele caused mild eye and head overgrowth (Figure 2A) (Tapon *et al*. 2002). Eye and head overgrowth was also observed when tissue was homozygous for the strong loss of function *CtBP* allele *CtBP^87De-10^*, consistent with prior studies (Hoang *et al*. 2010; Sumabat 2019), although the overgrowth phenotype was weaker than for *sav^2^* (Figure 2A). Strikingly, eye and head tissue that was double mutant for both *sav^2^* and *CtBP^87De-10^* displayed dramatic overgrowth, somewhat resembling null alleles of *sav* such as *sav^3^* and *sav^shrp1^* (Figure 2A) (Kango-Singh *et al*. 2002; Tapon *et al*. 2002). Similar results were obtained with the hypomorphic *CtBP^03463^* alelle, alone and in combination with *sav^2^* (Figure S1A). To assess these phenotypes in a more quantitative manner, we scored the number of interommatidial cells in the pupal eye 44 hrs after puparium formation, when developmental apoptosis has ceased in this tissue. An increase in interommatidial cells, is a feature of tissue mutant for Hippo pathway genes (Tapon *et al*. 2002). Pupal eyes harbouring the *CtBP^87De-10^* allele displayed normal interommatidial cell numbers (Figure 2A and B). As expected, *sav^2^* mutant eyes displayed a moderate increase in interommatidial cells, consistent with previous reports (Tapon *et al*. 2002). In *CtBP, sav* double mutant eyes, interommatidial cell number was almost doubled compared to *sav^2^* alone, consistent with the stronger overgrowth of double mutant tissue (Figure 2A and B).

**Figure 2.**
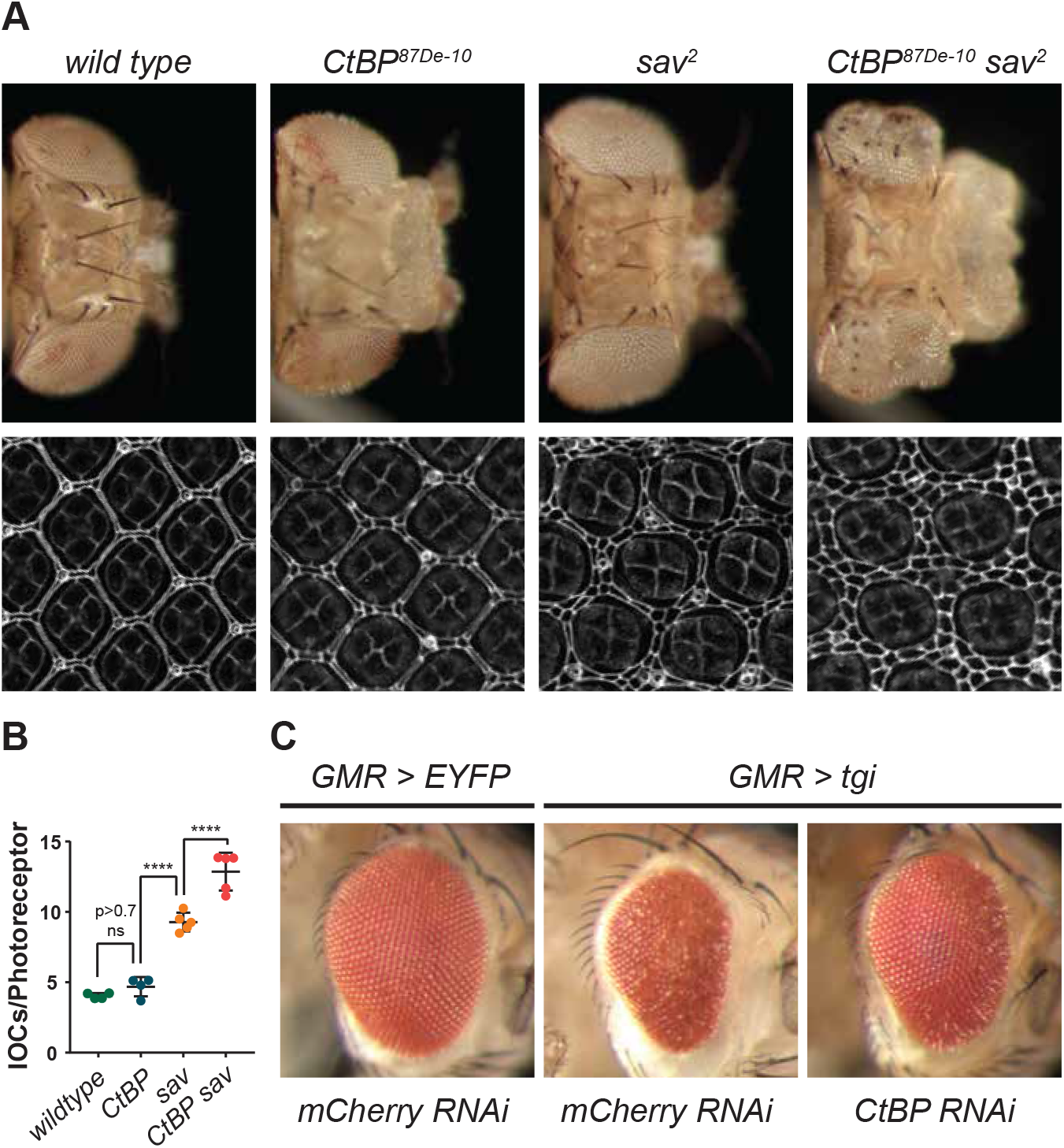
CtBP limits eye growth and is required for Tgi-induced growth inhibition. (A) Adult female *D. melanogaster* heads, anterior is to the right (upper panels) and pupal retinas 44 hr APF stained with Disc large (white) (lower panels). In each, tissues were predominantly comprised of cells that were homozygous for the indicated genotypes, which were generated by eyFLP-driven mitotic recombination over a cell lethal FRT82B chromosome. (B) Quantification of the number of interommatidial cells relative to photoreceptor cells in each ommatidial cluster. n = 4 for *wt* and *CtBP*, and 5 for *sav* and *sav, CtBP*. Data represent mean +/- standard deviation, **** p<0.0001 one way ANOVA with post hoc Tukey’s test ns, non-significant. (C) Adult male *D. melanogaster* heads from the indicated genotypes. *CtBP* RNAi line: *KK107313* 30B (details in Material and Methods). Anterior is to the right.

Next, we investigated genetic interactions between *tgi* and *CtBP*. Clones of tissue lacking *tgi* function in eye imaginal discs display no obvious growth phenotypes and do not cause deregulation of Yki/Sd target genes (Guo *et al*. 2013; Koontz *et al*. 2013). By contrast, *tgi* overexpression represses the growth of both the *D. melanogaster* eye and wing in a Sd-dependent manner (Guo *et al*. 2013; Koontz *et al*. 2013). Therefore, we investigated the functional requirement of *CtBP* for *tgi*-mediated growth suppression, using an eye-specific *tgi* overexpression system. *GMR-Gal4* driven expression of a *UAS-HA-tgi* transgene led to eye size reduction. Depletion of CtBP by RNAi strongly suppressed Tgi-induced growth retardation and this was observed using two independent CtBP RNAi lines (Figure 2C and S1B).

### Tgi requires Pits to suppress eye growth

Next, we addressed the requirement of Pits for Tgi-mediated growth suppression. Like CtBP, RNAi-mediated depletion of Pits suppressed eye growth inhibition by Tgi (Figure 3A). The Pits RNAi line we used was on target as it caused strong Pits knockdown, as assessed either by immunofluorescence or western blotting (Figure S2A and B). To further investigate a role for Pits, we used CRISPR-Cas9 genome editing to generate two different frameshift mutations in the *pits* gene (*pits^SK2^* and *pits^SK5^*), which both disrupted the *pits* open reading frame by generating frameshift mutations at amino acid 123 and subsequent stop codons (Figure S2C). Pits protein expression was undetectable in tissues harbouring these mutations, as determined by both western blotting and immunofluorescence (Figure S2D and E). As such, we predict that these are strong loss of function alleles in *pits*. In addition, we made use of a *pits* P element insertion strain (*pits^KG^*) that displayed reduced Pits protein expression (Figure S2C and D). All three *pits* alleles were homozygous viable and hemizygous viable and displayed no gross phenotypic abnormalities. This is consistent with a recent study that used imprecise P-element excision to generate three *pits* alleles, two of which were homozygous viable, whilst the third was semi-lethal (Liaw 2016). An independent study used recombination mediated cassette exchange to generate *pits* mutant alleles, which were hemizygous lethal (Marcogliese *et al*. 2018). We utilised our *pits* mutant alleles to further test whether Tgi requires *pits* to retard eye growth. Consistent with RNAi-mediated depletion of Pits, Tgi’s ability to repress eye growth was strongly suppressed in three independent strains of hemizygous *pits* animals (*pits^SK2^, pits^SK5^* and *pits^KG^*) (Figure 3A and S3F). This demonstrates that, like loss of *CtBP*, loss of *pits* suppresses Tgi-induced growth retardation in the *D. melanogaster* eye.

**Figure 3.**
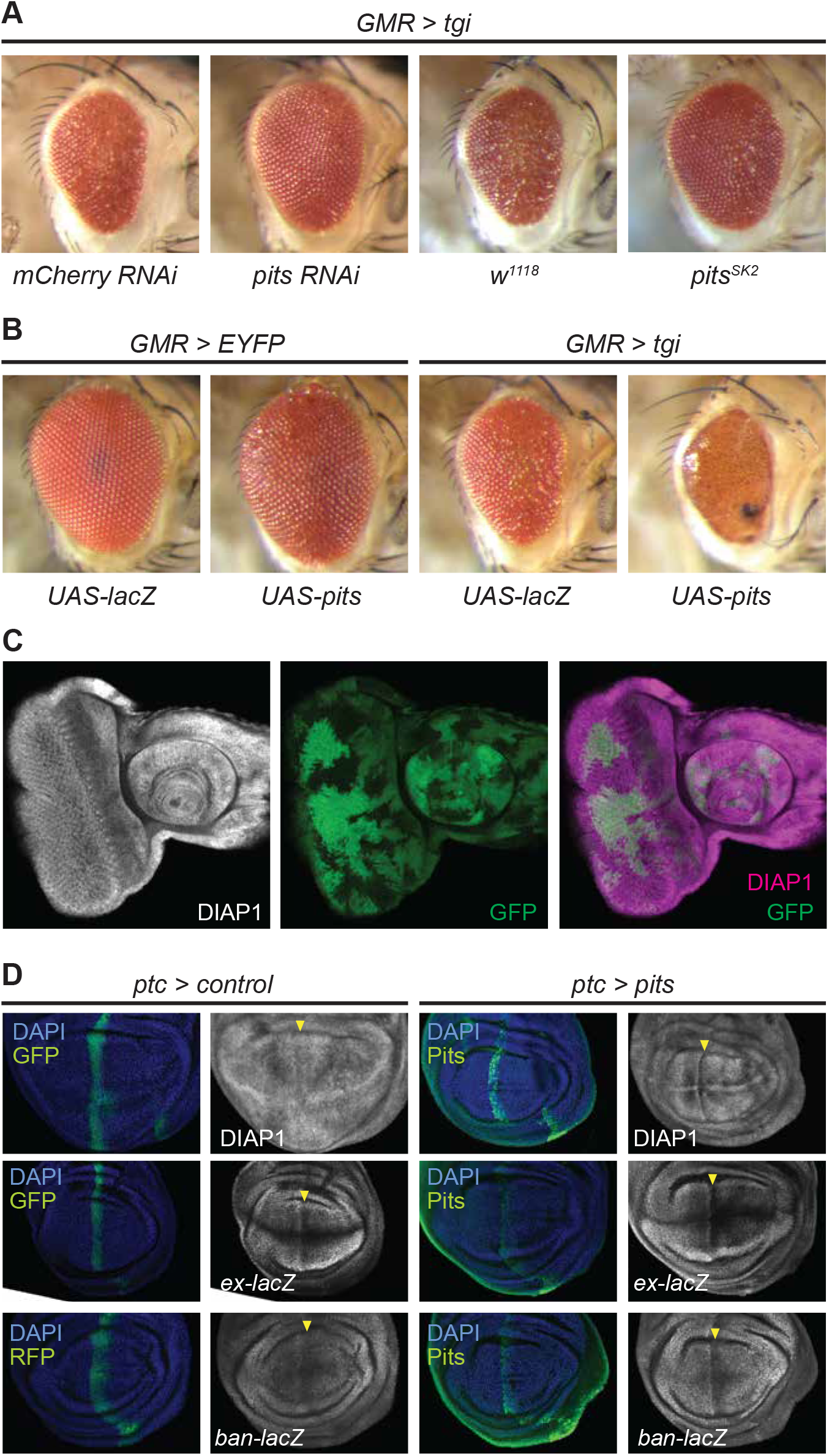
Pits is required by Tgi to suppress eye growth and can repress Hippo pathway target genes. (A and B) Adult male *D. melanogaster* heads from the indicated genotypes, anterior is to the right. (C) Third instar larval eye imaginal disc, anterior is to the right. GFP (green) marks wild-type tissue, whilst GFP-negative tissue harbours the *pits^SK2^* allele. DIAP1 protein is in greyscale in the left panel and magenta in the merged panel on the right. (D) Third instar larval wing imaginal discs of the indicated genotypes. DIAP1 protein, *ex-lacZ* and *ban-lacZ* are in greyscale. DAPI (blue) marks nuclei and GFP or RFP (green) marks the expression domain of the indicated transgenes. Yellow arrowheads indicate the expression domain of control or *pits* transgenes.

We also performed the converse experiment, where we employed *GMR-Gal4* to overexpress *pits* using *D. melanogaster* strains harbouring either a UAS-*pits* transgene that we generated (Figure 3B), or a P element inserted into the transcriptional start site of *pits* (Figure S3B). Overexpression of *pits* was confirmed in both strains by western blot (Figure S3A). *pits* overexpression alone had no effect on eye growth (Figure 3B; Figure S3B). However, when *pits* was overexpressed with *tgi* it enhanced *tgi*-mediated suppression of eye size (Figure 3B; Figure S3B). Moreover, partial lethality was observed in animals that overexpressed both *pits* and *tgi* in the eye, whereas expression of either transgene alone did not impact viability (Figure S3C).

### Pits overexpression can repress abundance of Hippo pathway target genes

Next, given the described roles of CtBP and Pits as transcription corepressors, we hypothesised that they play important roles in repression of Sd and Tgi target genes. Since CtBP is a general corepressor employed by a large number of transcription factors, we initially focussed on Pits. Previously, *tgi* overexpression was found to limit Yki/Sd target gene expression, whereas *tgi* loss of function had no impact on these genes (Guo *et al*. 2013; Koontz *et al*. 2013). Consistent, with these *tgi* loss of function studies, the abundance of the Yki/Sd target DIAP1 was unchanged in *pits* mutant eye imaginal disc clones (Figure 3C). We then switched to overexpression studies and found that *ptc-Gal4* driven overexpression of *tgi* suppressed expression of the Yki/Sd target DIAP1 in larval wing imaginal discs (Figure S3D), in accordance with previous studies (Guo *et al*. 2013; Koontz *et al*. 2013). Similarly, *ptc-Gal4* driven overexpression of *pits* caused downregulation of DIAP1 in larval wing imaginal discs (Figure 3D). To further investigate the impact of *pits* overexpression on well-established Yki/Sd target genes we utilised lacZ enhancer trap lines in *bantam (ban)* and *expanded (ex)*. *pits* overexpression also decreased abundance of both *ban-lacZ* and *ex-lacZ*, although to a lesser degree than DIAP1 (Figure 3D). These results show that Pits, like Tgi, can repress expression of well-known Yki/Sd target genes when overexpressed, but is not required to regulate expression of these genes in normally growing tissues.

### Loss of *CtBP*, but not *pits*, can suppress the undergrowth of *yorkie* mutant eye tissue

A role for Tgi in Sd-mediated transcriptional repression was revealed by generating larval eye discs that harboured both *yki* and *tgi* mutant clones. *tgi* loss partially suppressed the undergrowth of *yki* mutant clones, although it did not restore DIAP1 expression, which was reduced in *yki* clones (Koontz *et al*. 2013). Therefore, we performed similar experiments by generating eyes that were mosaic for both *yki* (RFP negative) and *ctbp* (GFP negative) mutant clones. Larval eye clones that were double mutant for both *yki* and *ctbp* mutant survived better than *yki* single mutant clones, however DIAP1 expression was not obviously restored (Figure 4A, B and E). This indicates that *ctbp* loss can partially restore the growth defects associated with *yki* loss. We also assessed a role for *pits* in Sd-mediated repression of transcription and eye growth by generating *yki* mosaic eye discs in *pits* homozygous mutant animals or wild-type control animals. The size of *yki* mutant clones did not obviously differ when they were generated in a wild-type background or *pits* mutant background, however (Figure 4C, D and F). This shows that *pits* is not an essential mediator of Sd-mediated transcriptional repression, at least in the conditions tested here.

**Figure 4.**
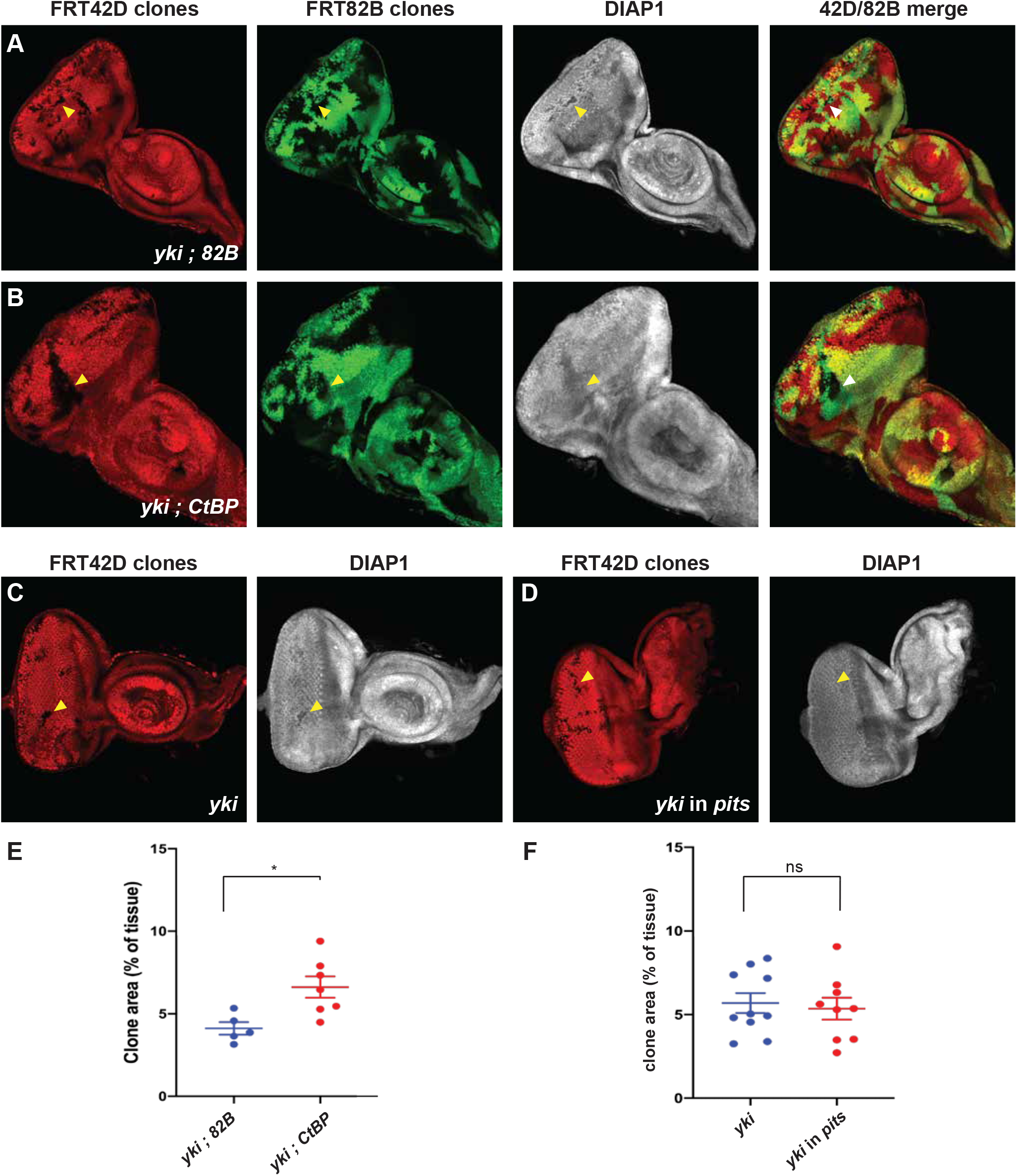
Loss of *CtBP*, but not *pits*, can suppress the undergrowth of *yorkie* mutant eye tissue. (A-D) Third instar larval eye imaginal discs, anterior is to the right. RFP (red) marks tissue wildtype for FRT42D, whilst RFP negative tissue is mutant for *yki*. DIAP1 protein expression is in greyscale. In (A) both GFP positive (green) and negative tissue are wildtype for FRT82B. In (B) GFP negative tissue is mutant for *CtBP*. In (A) and (B) merged images of GFP and RFP are shown on the far right. In (C) *yki* mutant clones were generated in a wildtype background whilst in (D) they were generated in a *pits* mutant background. Select *yki* mutant clones are indicated with yellow arrowheads. (E and F) Chart showing quantification of yki mutant clone area expressed as a % of the whole tissue. In (E) clones possessed either a *yki* mutant and wildtype FRT82B chromosome, or a *yki* mutant and *CtBP* mutant chromosome. In (F) *yki* mutant clones were generated in either wildtype animals or *pits* mutant animals. n = 5 and 7, respectively in (E) and n = 10 and 9, respectively in (F). Data represent mean +/- SEM, * p<0.05 t-test. ns, non-significant.

## DISCUSSION

The transcriptional corepressor Tgi has emerged as an important regulator of transcription that is regulated by Yki and Sd but the mechanism by which it does so is currently unclear (Guo *et al*. 2013; Koontz *et al*. 2013). We set out to address this by identifying Tgi-interacting proteins. Four such proteins were identified with high confidence, all of which have been ascribed functions as transcription regulators: the previously identified Hippo pathway proteins Yki and Sd, as well as Pits and CtBP. Previous structure-function studies showed that Tgi’s growth inhibitory function fully depends on its ability to interact with Sd (Guo *et al*. 2013; Koontz *et al*. 2013), suggesting that Sd is the sole transcription factor that mediates Tgi’s influence on transcription. Our finding that Sd was the only sequence-specific TF detected by mass spectrometry in Tgi purifications in S2 cells, further supports this model. Additionally, our biochemical data support the notion that Pits and CtBP function together with the Tgi corepressor to limit tissue growth, although our genetic studies fall short of providing conclusive evidence for this. The fact that *pits* and *ctbp* were required for Tgi’s ability to limit eye growth when overexpressed argues that Tgi requires them to repress gene expression through Sd. Further support for this idea comes from the finding that *pits* overexpression repressed the expression of well-defined Yki/Sd/Tgi target genes like *DIAP1*, *ban* and *ex*. On the other hand, *pits* mutant flies were homozygous viable and displayed no obvious gross phenotypic abnormalities, and *pits* mutant larval eye imaginal discs cells expressed normal levels of DIAP1. However, it should be noted that loss of either *sd* or *tgi* in larval eye imaginal discs also has no obvious impact on expression of its target genes, whilst overexpression does (Guo *et al*. 2013; Koontz *et al*. 2013). This suggests that in the growing larval eye imaginal disc Sd does not have a major role in gene repression. Alternatively, in the absence of *sd, tgi* or *pits*, other proteins might compensate for them and regulate expression of their target genes.

Strong genetic evidence that helped identify Tgi as a mediator of Sd’s transcriptional repression activity was the finding that loss of *tgi* partially rescued the undergrowth of *yki* clones in larval eye imaginal discs (Koontz *et al*. 2013). Similarly, in our study, loss of *CtBP* partially restored the undergrowth phenotype of *yki* larval eye clones. Therefore, CtBP might work in partnership with Tgi and Sd to repress target genes that are required for eye growth. Alternatively, CtBP might limit eye overgrowth by acting in parallel to Tgi and possibly also Sd. Indeed, a recent study provided evidence for both of these models with the discovery that CtBP represses transcription of the progrowth microRNA *bantam* in both Yki-dependent and independent manners (Sumabat 2019). In contrast to *CtBP* loss, *pits* loss did not rescue the undergrowth phenotype of *yki* eye imaginal disc clones, which stands in apparent opposition to its requirement for growth inhibition induced by Tgi overexpression. The reason for this discrepancy is currently unclear. Future studies aimed at identifying the full suite of Yki/Sd/Tgi target genes that are required for mediating eye growth should help to provide clarity on the different roles of Yki, Sd, Tgi, CtBP and Pits on transcription and eye growth, but this genetic program is currently unknown.

The mechanism by which Yki/Sd/Tgi regulate transcription in *D. melanogaster* appears to be largely conserved in mammals (Guo *et al*. 2013; Koontz *et al*. 2013; Zhang *et al*. 2014). Here, we found that Pits and CtBP bind to Tgi through conserved protein motifs. Therefore, their biochemical relationship is also likely to be conserved in other species. Indeed, the interaction between the human CtBP and Tgi orthologues, CTBP2 and VGLL4, respectively has been reported, which inhibits adipogenesis of murine 3T3-L1 cells (Zhang *et al*. 2018). In addition, several studies have identified a physical interaction between VGLL4 and the human Pits orthologues, IRF2BP1, 2 and L (Teng *et al*. 2010; Huttlin *et al*. 2015). A very recent study reported a functional interaction between these genes in the context of cancer; the Pits orthologue IRF2BP2 acted with VGLL4 to suppress liver tumor growth that was caused by YAP hyperactivity and also the expression of YAP-TEAD target genes (Feng *et al*. 2019). Our *D. melanogaster* studies imply that Pits can act as a corepressor of Yki/Sd target genes, as opposed to an activator. In support of this, most studies of the three mammalian Pits orthologues, IRF2BP1, 2 and L, indicate that they act as corepressors (Stadhouders *et al*. 2015; Liaw 2016; Ramalho-Oliveira *et al*. 2019), although the mechanism of co-repression is poorly characterized, and CtBP is generally considered a transcription repressor (Turner and Crossley 2001). Clearly, further studies are required to clarify the mechanism by which Hippo pathway target genes are regulated by Yki, Sd and Tgi. To explore roles for Pits and CtBP on transcription in the growing eye it will be important to examine their genome occupancy relative to Tgi and Sd, and also to better define the mechanism by which they regulate transcription. This should shed light on the control of Hippo pathway target genes and could also be valuable for defining the emerging role of the Tgi orthologue VGLL4 in human cancers.

## ACKNOWLEDGEMENTS

We thank V. Corces, D. Pan, N. Tapon, A. Veraksa, the Vienna *D. melanogaster* RNAi Center, the Australian *Drosophila* Research Support Facility (www.ozdros.com), the Bloomington *Drosophila* Stock Center and the Developmental Studies Hybridoma Bank for *D. melanogaster* stocks, plasmids and antibodies. We thank I. Hariharan for discussions of unpublished data. We acknowledge the Peter Mac Microscopy and Histology facility and the University of Melbourne Bio21 Mass spectrometry facility. K.F.H is a National Health and Medical Research Council Senior Research Fellow (APP1078220). J. V. was supported by a Victorian Cancer Agency Early Career Seed Grant (ECSG14026). This research was supported by the National Health and Medical Research Council of Australia (APP1080131).

---

**Supplementary Figure 1.**
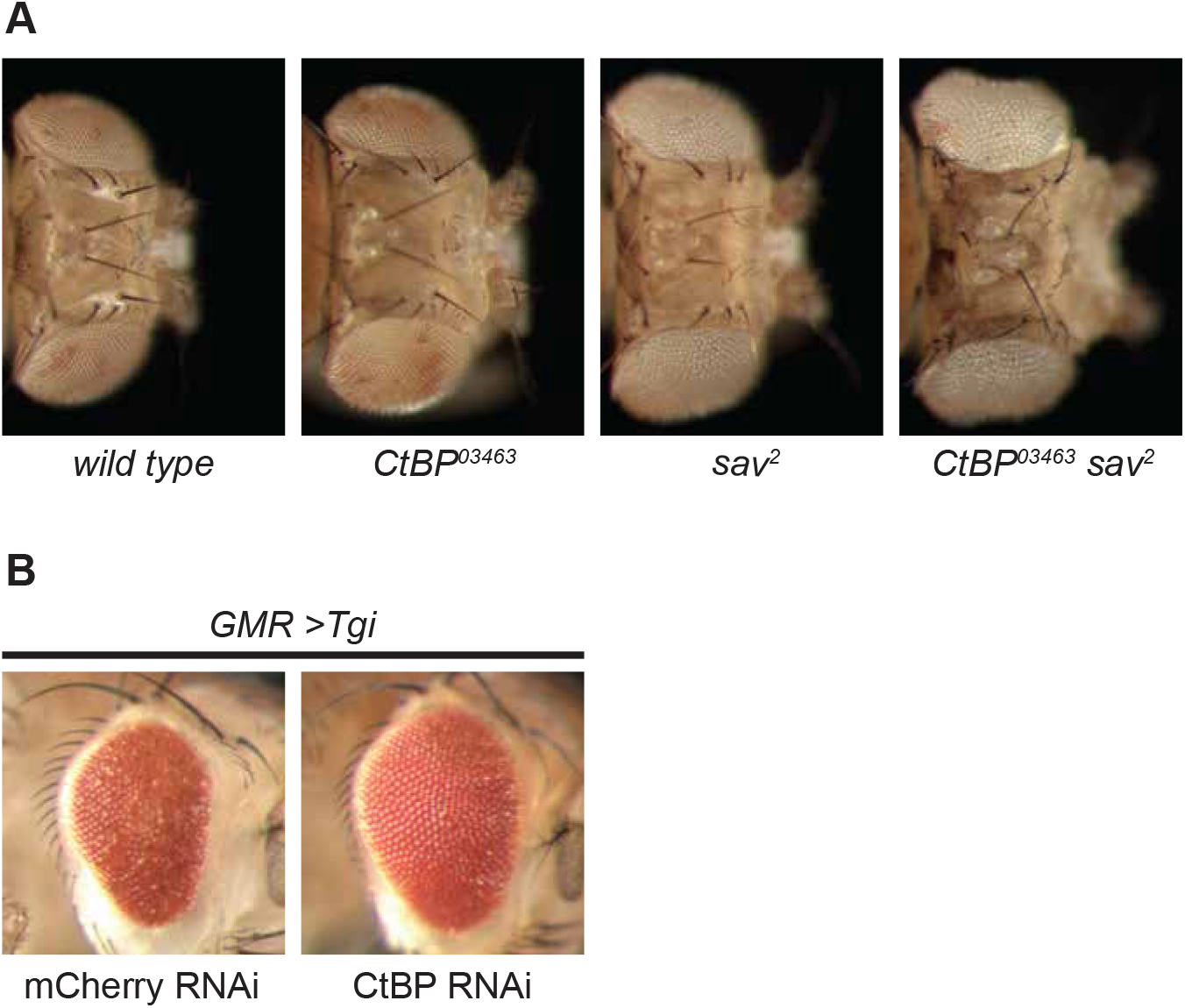
CtBP limits eye growth and is required for Tgi-induced growth inhibition. (A) Adult female *D. melanogaster* heads, anterior is to the right. Tissues were predominantly comprised of cells that were homozygous for the indicated genotypes, which were generated by eyFLP-driven mitotic recombination over a FRT82B cell lethal chromosome. (B) Adult male *D. melanogaster* heads from the indicated genotypes. CtBP RNAi line: *BSC#32889*. Anterior is to the right.

**Supplementary Figure 2.**
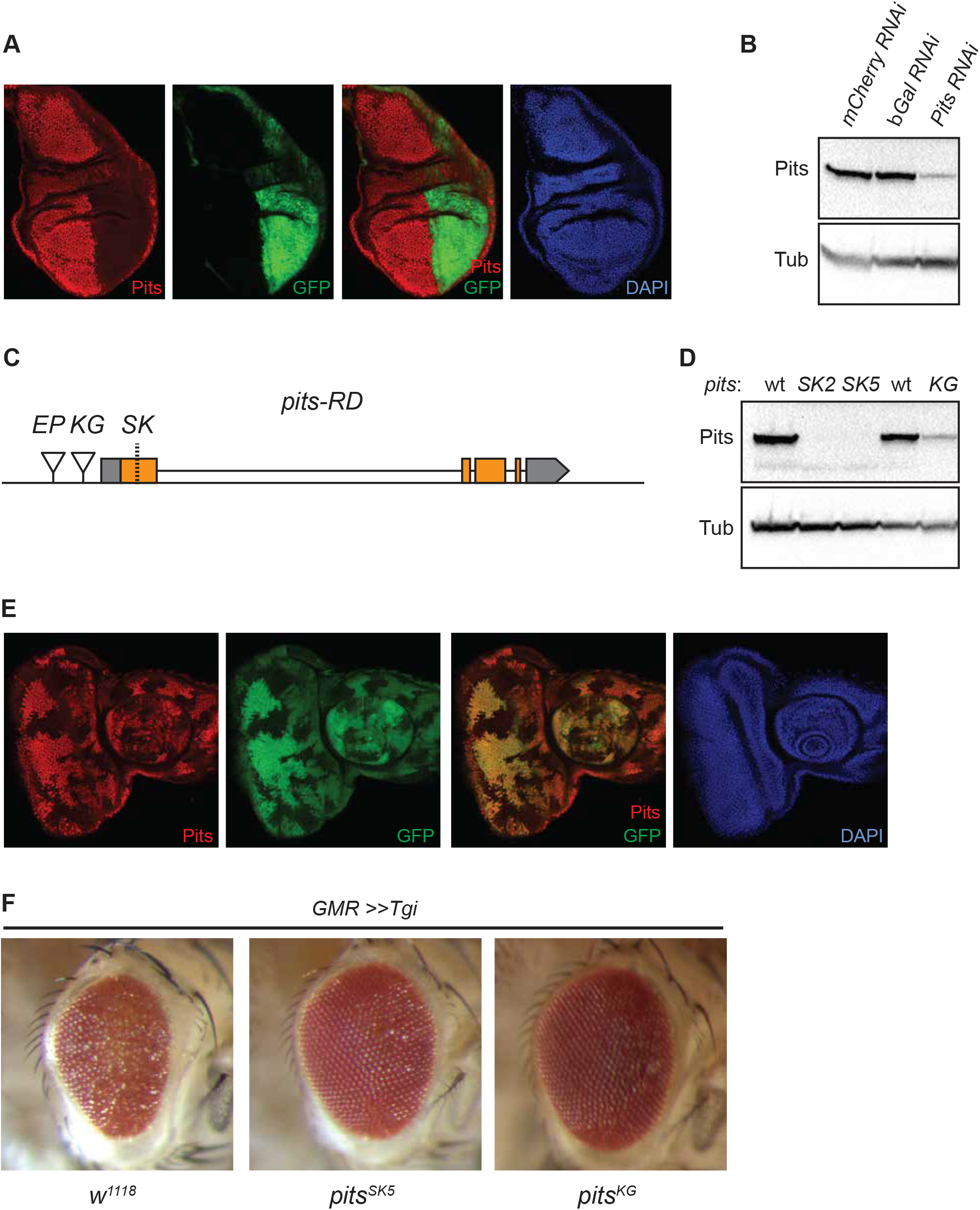
Pits is required by Tgi suppress eye growth. (A) A third instar larval wing imaginal disc that expresses a Pits RNAi transgene in the posterior compartment under the control of en-Gal4. Pits expression is in red, GFP (green) marks the posterior compartment. The third panel is a merge of Pits and GFP, whilst DAPI (blue) marks nuclei in the panel on the right. (B) Heads of adult flies harbouring tubulin-Gal4 crossed to the indicated RNAi transgenes were subjected to western blot analysis using the indicated antibodies. (C) Schematic diagram of the *pits* locus. Locations of P element insertions *P(EP)PITS^EP1313^* (EP), *P(SUPor-P)PITS^KG07818^* (KG) and CRISPR/Cas9-generated frameshift mutations (SK) are indicated relative to *pits* transcript variant D, which is the longest ORF of three *pits* transcript variants. (D) Heads of adult flies of the indicated genotypes were subjected to western blot analysis using the indicated antibodies. (E) The same third instar larval eye imaginal disc as in Figure 3D, anterior is to the right. Pits expression is in red, GFP (green) marks wild-type tissue, whilst GFP-negative tissue harbours the *pits^SK2^* allele. The third panel is a merge of Pits and GFP, whilst DAPI (blue) marks nuclei in the panel on the right. (F) Adult male *D. melanogaster* heads, anterior is to the right. Flies expressed *GMR-Gal4* and *UAS-HA-Tgi* and were hemizygous for either *w^1118^* (control), *pits^SK5^* or *pits^KG07818^*.

**Supplementary Figure 3.**
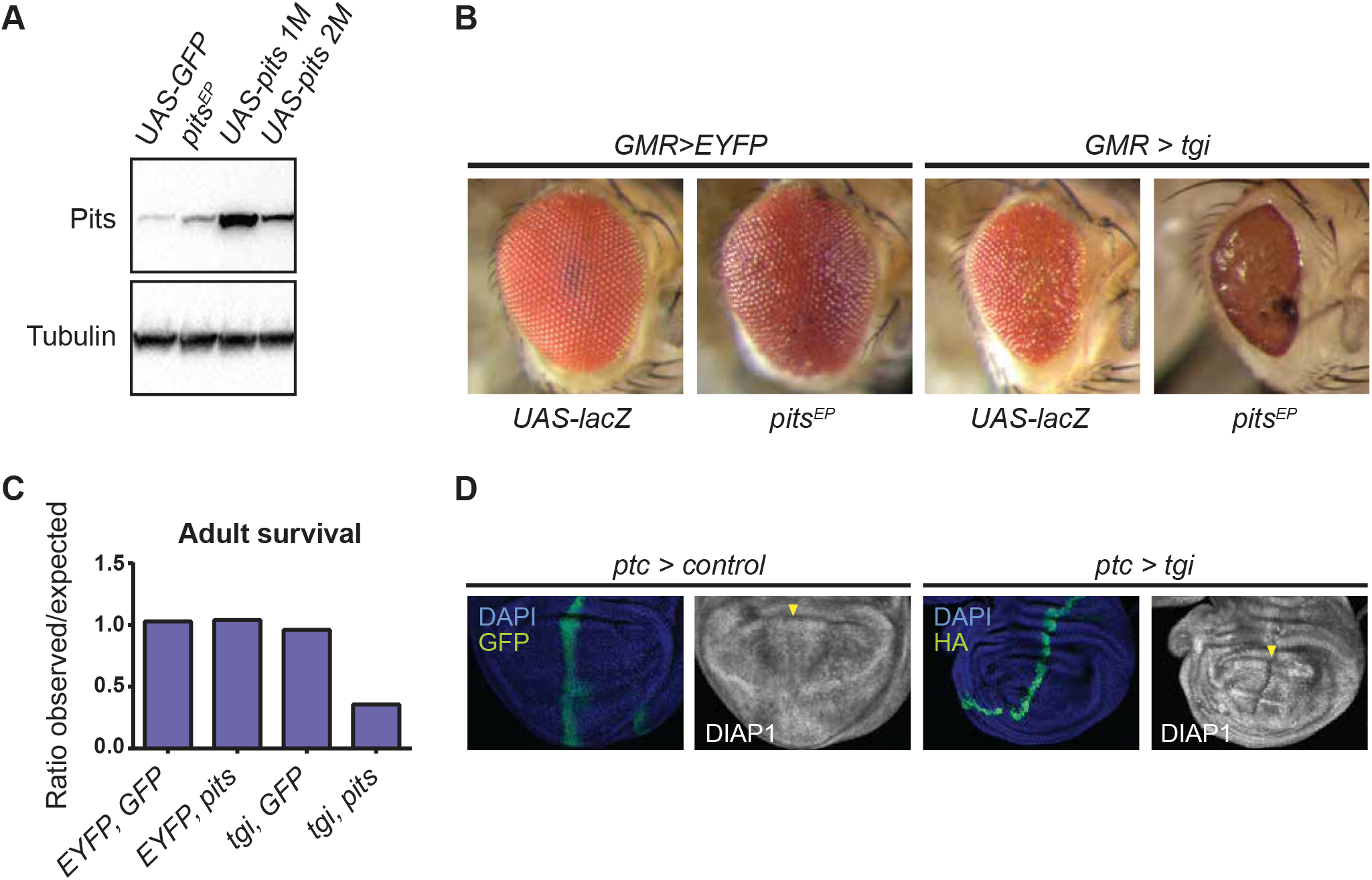
Cooperative effects of Tgi and Pits overexpression on eye growth and *D. melanogaster* survival. (A) Heads of adult flies harbouring GMR-Gal4 crossed to the indicated transgenes were subjected to western blot analysis using the indicated antibodies. (B) Adult male *D. melanogaster* heads, anterior is to the right. Flies expressed GMR-Gal4 and either *UAS-EYFP* or *UAS-HA-Tgi*, and either *UAS-lacZ* (control) or *pits^EP1313^*. (C) Relative amounts of viable adult *D. melanogaster* after *GMR-Gal4* driven expression of the indicated transgenes. The numbers of *D. melanogaster* counted were (relevant genotype/total): 157/305 (*EYFP, GFP*), 258/496 (*EYFP, pits*), 161/335 (*Tgi, GFP*), 45/505 (*Tgi, pits*). (D) Third instar larval wing imaginal discs of the indicated genotypes. DIAP1 protein is in greyscale. DAPI (blue) marks nuclei and GFP (green) marks the expression domain of the indicated transgenes. Yellow arrowheads indicate the expression domain of control or *pits* transgenes. Note the same control tissue is displayed here as in the related figure, Figure 3D.

